# Sterol derivative binding to the orthosteric site causes conformational changes in an invertebrate Cys-loop receptor

**DOI:** 10.1101/2023.01.06.523009

**Authors:** Steven De Gieter, Eveline Wijckmans, Diletta Pasini, Chris Ulens, Rouslan G. Efremov

## Abstract

Cys-loop receptors or pentameric ligand-gated ion channels are mediators of electrochemical signaling throughout the animal kingdom. Because of their critical function in neurotransmission and high potential as drug targets, Cys-loop receptors from humans and closely related organisms have been thoroughly investigated, whereas molecular mechanisms of neurotransmission in invertebrates are less understood. When compared with vertebrates, the invertebrate genomes underwent a drastic expansion in the number of the nACh-like genes associated with receptors of unknown function. Understanding this diversity contributes to better insight into the evolution and possible functional divergence of these receptors. In this work, we studied orphan receptor Alpo4 from an extreme thermophile worm *Alvinella pompejana*. Sequence analysis points towards its remote relation to characterized nACh receptors. We solved the first cryo-EM structure of a lophotrochozoan nACh-like receptor in which a CHAPS molecule is tightly bound to the orthosteric site. We show that the binding of CHAPS leads to extending of the loop C at the orthosteric site and a clockwise quaternary twist between extracellular and transmembrane domains. Both the ligand binding site and the channel pore reveal unique features. These include a conserved Trp residue in loop B of the ligand binding site which is flipped into an apparent self-liganded state in the apo structure. The ion pore of Alpo4 is tightly constricted by a ring of methionines near the extracellular entryway of the channel pore. Our data provide a structural basis for a functional understanding of Alpo4 and hints towards new strategies for designing specific channel modulators.

## Introduction

Cys-loop receptors are allosterically regulated pentameric ligand-gated ion channels (pLGICs). Functionally pLGICs are classified into cation- and anion-selective receptors. The former class is exemplified by nicotinic acetylcholine (nAChRs) and serotonin (5-HT_3_) receptors^1^. Anion-selective receptors are comprised of glycine (GlyRs) and GABA_A_ receptors. Because of their role in synaptic transmission, the inflammatory response and implication in diseases including gastro-intestinal, psychiatric and cognitive disorders, startle disease, epilepsy, and smoking addiction^2–5^, human and mammalian homologs of pLGICs have been the subject of intense research.

Recent advances in cryo-EM led to a rapid expansion of the structural data on nAChRs. The structures of 3 types of heteropentameric nicotinic receptors (α4β2, α3β4 and *Torpedo* muscle-nAChR) and one homopentameric receptor (α7) were determined^6–10^. For the α7 nicotinic receptor, structures of three major conformational states were solved: a resting state, an agonist-bound activated, and an agonist-bound desensitized state. Cryo-EM structures also have been determined for 5-HT_3_ receptors^11–13^, glycine receptors^14,15^ and GABA_A_ receptors^16–20^. Currently, only one structure of a non-vertebrate ion channel, the glutamate-gated chloride channel (GluCl), from *C. elegans* has been solved. GluCl is the drug target for anthelmintics such as ivermectin highlighting the importance of structural characterization of non-vertebrate pLGICs^21,22^.

Lophotrochozoa comprises one of the largest groups in the animal kingdom and includes organisms such as annelids, mollusks, and platyhelminths (flatworms). Intriguingly, genome analysis revealed a massive expansion of nAChR genes in these organisms with 52 and 217 nAChR genes identified in mollusks and annelids, respectively ^23^. This contrasts with the number of receptors found in organisms having an advanced nervous system, in which only 10 to 20 nAChRs are encoded in the genomes, for example 17 in humans^7^. It has been speculated that the expansion in nAChR genes is a consequence of the adaptation to a stationary lifestyle in a dynamic environment^23^. The biological role of the additional nAChRs and their properties remain unknown. Characterization of these receptors may lead to discoveries of alternative neurotransmitters, new signaling pathways as well as a better understanding of the evolution of neurotransmission.

We have previously biochemically characterized seven invertebrate Cys-loop receptors, Alpo1-7, identified in the proteome of *Alvinella pompejana*, an annelid worm that inhabits the surroundings of hydrothermal vents and is the most extreme thermophilic invertebrate currently known^24,25^. Among seven Alpo receptors, we identified two nAChR-like receptors (Alpo1 and Alpo4) and one Gly-like receptor (Alpo6), which were expressed and purified in amounts suitable for structural studies^26^. Alpo4 has 27-29% sequence identity with α-subunits of nAChRs and 25% with 5-HT_3_ receptors. The high biochemical stability and preliminary characterization using negative stain electron microscopy suggested that Alpo4 was a promising target for structural studies^27^. However, despite exhaustive screening in different expression systems in combination with a compound library, including acetylcholine and serotonin, we could not identify the agonist/neurotransmitter for Alpo4, thereby limiting functional studies^26^. Because of its biochemical stability and unique position between nAChRs and 5-HT_3_ receptors we characterized the structure of Alpo4 using single-particle electron cryogenic microscopy (cryo-EM).

## Results

### Alpo4 is an isolated member of lophotrochozoan nAChRs

A massive expansion of nAChR genes in lophotrochozoans suggests their importance in functional diversity and adaptations. In *A. pompejana* the total number of nAChR genes is not known because its genome has not been fully sequenced. To get further insight into the relation of Alpo4 to other nAChRs we applied comparative genomic analysis. We performed a phylogenetic comparison of Alpo1-4 with nAChRs from annelids: *Capitella teleta (CT), Dimorphilus gyrociliatus* (DM), *Owenia fusiformis* (OW), *Hirudo verbana* (HV), *Helobdella robusta* (HR), and from mollusca: *Crassostrea virginca* (MV), *Crassostrea gigas* (CG), *Mizuhopecten yessoensis* (MY), *Pecten maximus* (PM) and *Pomacea canaliculata* (PC)^27^. A total of 649 sequences were grouped into 25 families (Supplementary Figure 1a). In the phylogenetic tree some sequences cluster in molluscan-specific families (e.g., 58A, 58B, 61A, 62A, and 63A), whereas other families, like 33A, includes members from all lophotrochozoan and has characteristic features of a vertebrate α7 subunit^28^. Alpo1-4 were classified into different sequence families. Alpo2 and 3 are found in the families 32C and 41A (sequence identities of the closest homolog are 45% and 65%, respectively), both of which contain sequences from the genome of each included lophotrochozoan suggesting functional importance and conservation of these protein families within the clade (Supplementary Figure S1b). Interestingly, in Polychaeta organisms, the characteristic vicinal disulfide in the tip of loop C required for ligand binding is not present in Alpo3-like nAChRs.

On the contrary, Alpo1 and Alpo4 (families 42A and 43A; sequence identities of the closest homolog are 36% and 37%) do not clusters in the well-populated families (Supplementary Figure 1b). This suggests either a unique function or a faster evolution. To further characterize Alpo4, we proceeded with its structural characterization by cryo-EM.

### Structure of Alpo4 reveals CHAPS bound to the orthosteric site

We purified Alpo4 in the detergent LMNG following the protocol established in Wijckmans 2016 et al. and used cryo-EM to solve its structure. The map reconstructed to 4.1 Å resolution confirmed that Alpo4 assembles into a homopentamer and has a conserved architecture of the pLGIC family (Figure 1a, Table 1). Each Alpo4 subunit is composed of a β-sandwich extracellular domain (ECD) and a transmembrane domain (TMD) made of four trans-membrane α-helices M1-M4 (Figure 1b, Supplementary Figure 2). Helices M2 contributed by each subunit are radially arranged around a central ion-conducting pore. The density for the intracellular domain (ICD), residues 308-412, was missing, and it was consequently not modeled (Supplementary Figure 3). An additional density next to the side chain of N167 on ECD was modeled as a N-acetylglucosamine (GlcNAc; Figure 1b, Supplementary Figure 2b). Glutamine glycosylation at the structurally equivalent position was also found in the 5-HT_3_ receptor, but not in nicotinic receptors^29^.

**Figure 1.**
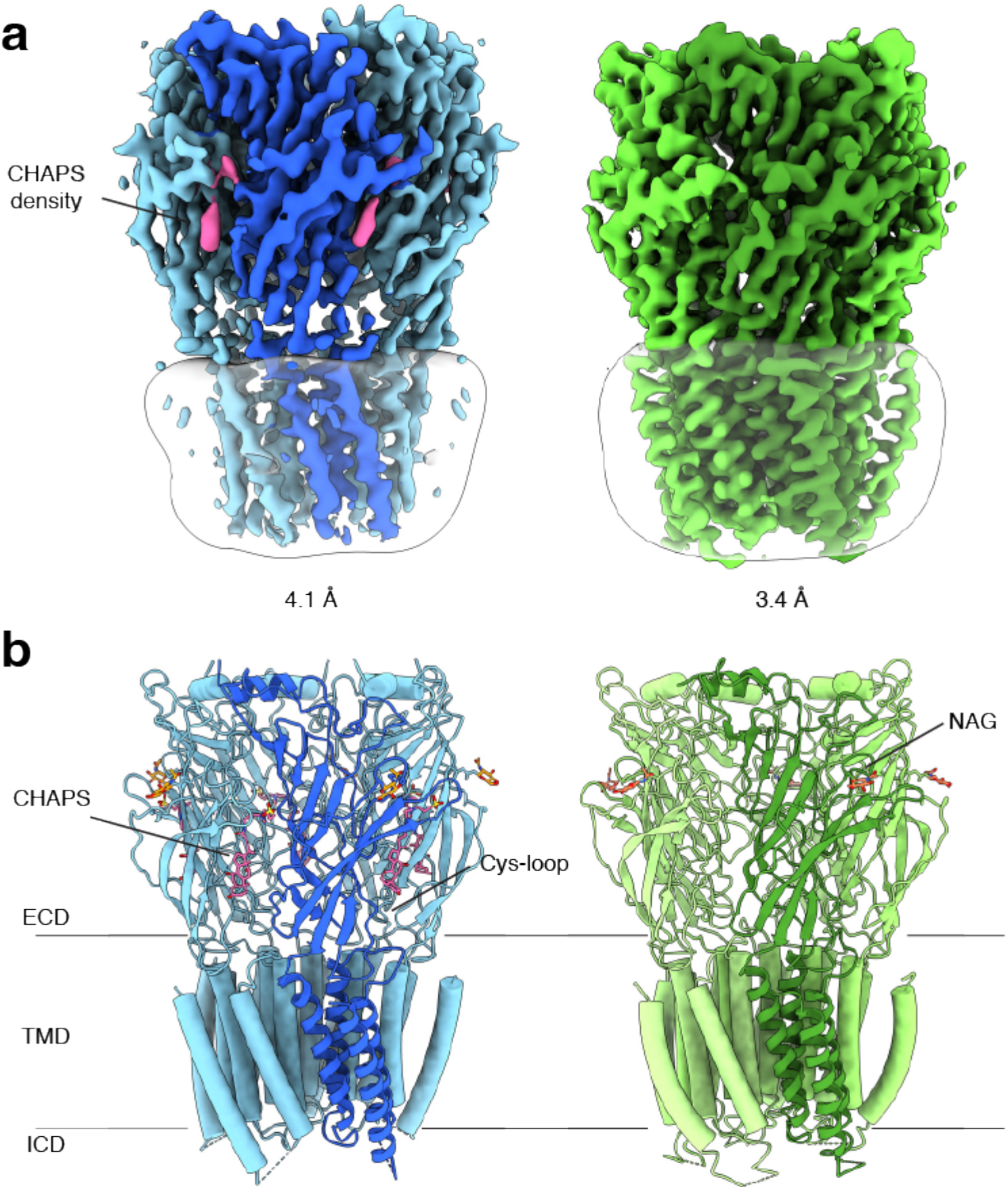
Architecture of Alpo4 in detergent. **(a)** Cryo-EM reconstruction of apo Alpo4^CHAPS^ (blue) and Alpo4^APO^ (green) at a resolution of 4.1 Å and 3.4 Å, respectively. The detergent micelle is shown in white surface representation. A monomer is shown in a darker shade. The density corresponding to bound CHAPS is shown in violet. **(b)** Side view of the atomic models shown in cartoon representation with NAG moieties shown as sticks. One subunit is highlighted. Bound CHAPS molecules are shown as sticks (violet).

Although no ligand was added, an additional density was observed in the orthosteric binding site located at the interface between the ECDs (Figure 1a, Supplementary Figure 4a). Despite the limited resolution, this density was well-resolved and consistent with a CHAPS molecule, a steroid-derived detergent. We further refer to this structure as Alpo4^CHAPS^. CHAPS was present in the purified Alpo4 at a concentration of 0.007% (110 μM) because of its thermostabilizing effect on Alpo4^26^. At 110 μM, the CHAPS concentration is 70-fold lower than the critical micellar concentration (CMC) of the detergent, suggesting a specific interaction with Alpo4 beyond modulating properties of the detergent belt and therefore is consistent with CHAPS binding to the orthosteric site^31^.

### Structure of the ECD and ligand-binding pocket with CHAPS

The ECD comprises an amino-terminal α-helix followed by ten β-strands folded into a β-sandwich (Figure 1b, Supplementary Figure 2b). CHAPS is bound at the ligand-binding pocket located at the interface of the principal (loops A-C) and the complementary subunit (loops D-F; Figure 2a, d). The CHAPS-Alpo4 interactions can be divided into two regions: the hydrophilic moiety and the sterol-binding moiety (Supplementary Figure 4a). The hydrophilic moiety, in part formed by the dimethylammonio group, shares structural resemblance with carbachol and overlaps with the canonical Cys-loop receptor ligand-binding site, involving a group of highly conserved aromatic residues F103 (loop A), W159 (loop B), Y199 and Y205 (loop C) of the principal subunit and W65 (Loop D) of the complimentary subunit. Here, the quaternary ammonium group of the CHAPS molecule establishes cation-π interaction with W159 (Figure 2a, d). This interaction is strikingly similar to the cation-π interactions observed with the quaternary ammonium group of the carbachol-bound AChBP structure (PDB: 1UV6), or the pyrrolidine nitrogen group in the nicotine-bound α4β2 nAChR structure (PDB: 5KXI) (Figure 2b, e). The interaction with the hydrophilic moiety is further stabilized by a salt bridge between the sulfonate group of CHAPS and Alpo4-specific Arg171 (Figure 2a).

**Figure 2.**
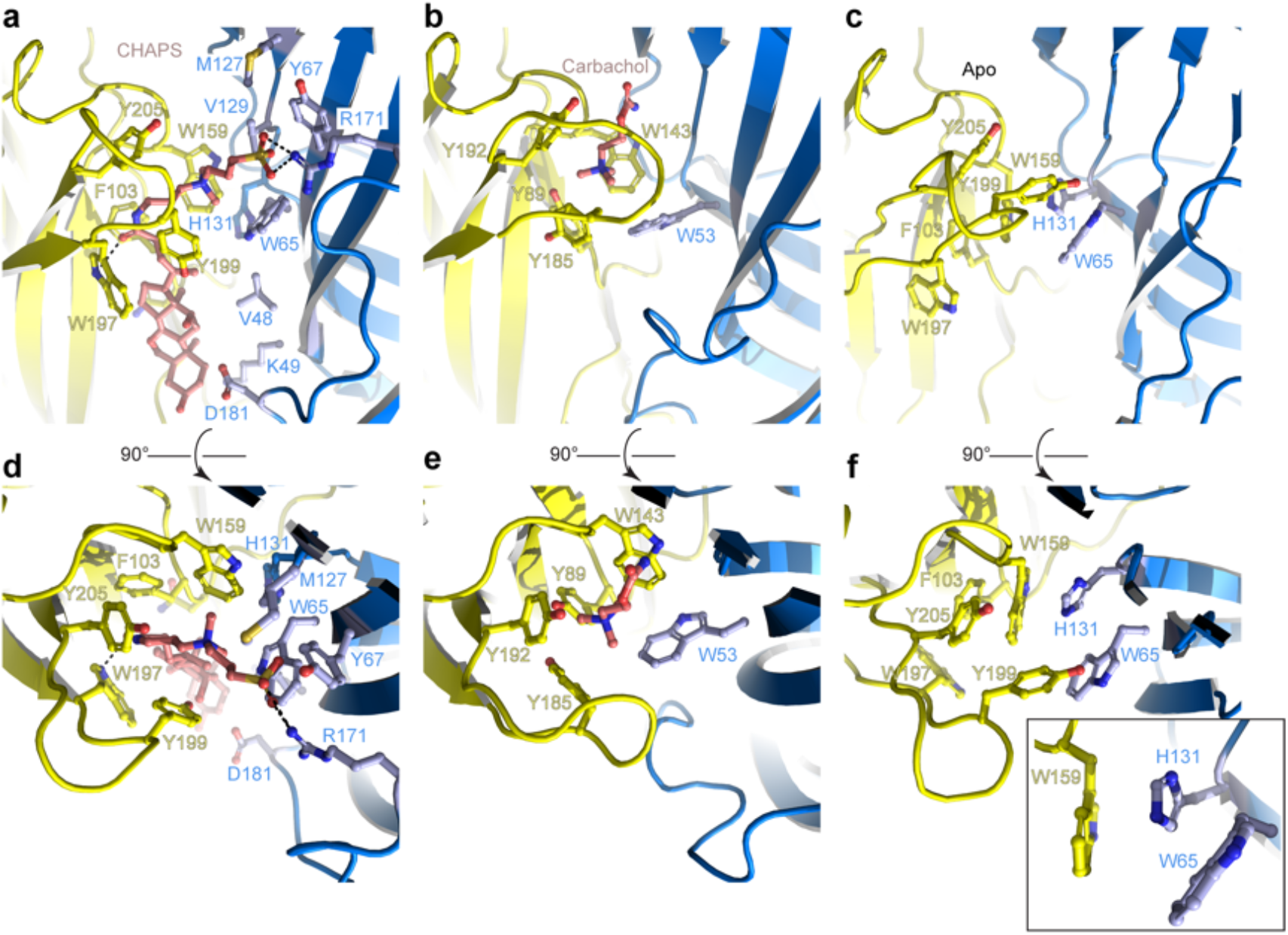
Orthosteric binding site of Alpo4. **(a, d)** Ligand binding pocket of Alpo4 with bound CHAPS. The residues interacting with CHAPS are shown as sticks. The principal subunit is colored yellow and the complementary is blue. **(b,e)** Similar views of the ligand binding site in the acetylcholine binding protein (AChBP) in complex with carbachol (PDB code 1UV6). **(c,f)** Ligand binding pocket of Alpo4 in apo state. Residues constituting the aromatic cage are shown as sticks. The self-liganded state of W159 is shown as an inset in sticks representation.

The sterol ring moiety of CHAPS is composed of three cyclohexane and one cyclopentane rings and it fits into a hydrophobic crevice with a high shape-complementarity. The sterol ring interacts with residues F103 (loop A), N104, F137, V155, W197 on the principal side and residues V48, K49, and D181 on the complementary side exclusively via Van der Waals contacts (Figure 2a, Supplementary Figure 4). This hydrophobic crevice is specific to Alpo4 and it is lined by poorly conserved residues. In other nAChRs, the pocket is narrower (Supplementary Figure 4b) and is lined by multiple charged residues. These molecular interactions of CHAPS with Alpo4 explain why the binding of the detergent is specific.

### Structure of the ligand-binding pocket in the apo state

To gain further insight into the structural and ligand-binding properties of Alpo4, several structures were solved in the absence of CHAPS. This was accomplished in two ways. Either CHAPS was removed from the Alpo4^CHAPS^ using size exclusion chromatography or we purified Alpo4 without any CHAPS present (Table 1). The highest resolution reconstruction of 3.4 Å was obtained from the protein sample that was depleted of CHAPS using size-exclusion chromatography (Supplementary Figure 5). This reconstruction, which we refer to as the apo state (Alpo4^APO^), allowed detailed modeling of the atomic structure of Alpo4 in most parts of the density (Figure 1b). Despite the higher overall resolution of the reconstruction, the density corresponding to the tip of the C- and the F-loops were less well resolved which indicates their increased flexibility in the absence of a ligand. At low sigma levels, residual densities were observed in the ligandbinding and the sterol-binding pockets suggesting that residual CHAPS was bound at low occupancy.

Similar results were obtained from the sample prepared in LMNG without CHAPS, reconstructed to 4.1 Å (Supplementary Figure 7). This structure is essentially identical to the apo state with (RMSD of 0.9 Å) (Supplementary Figure 8). No residual density was observed in the ligand-binding pocket in the later reconstruction, supporting the assignment of the 3.4 Å map as an apo state. The conformation of the aromatic residues constituting the ligand-binding pocket differed from the CHAPS-bound state. Specifically, W159 (loop B) flips and forms a cation-π interaction with H131, while being pinched by W65 (loop D) (Figure 2f). In this conformation, W159 can no longer form a cation-π interaction with an external ligand, therefore it is tempting to speculate that this conformation represents a “self-liganded” state.

Given that Alpo4 shares a structural resemblance to the α4β2 nAChR and the 5-HT_3_R, we investigated whether acetylcholine or serotonin binds into the ligandbinding site. To this end, structures of Alpo4 purified in the absence of CHAPS and with an added, 1 mM ACh or 1 mM serotonin were solved to a resolution of 3.9 Å and 6.2 Å, respectively (Supplementary Figures 7 and 9). Their overall conformation was identical to that of the apo state with an overall RMSD of 0.87 and 0.88 Å, respectively. No density in the ligand-binding site was observed in the reconstructions. Although 6.2 Å resolution is too low to interpret the density of serotonin, the overall quaternary structure was identical to that of the apo state. The absence of a quaternary twist expected for a desensitized conformation suggests that serotonin was not bound.

The cryo-EM reconstructions of Alpo4 obtained in the presence of acetylcholine and serotonin suggest that neither of the neurotransmitters binds Alpo4. This agrees with our electrophysiological experiments in various expression systems (*Xenopus* oocytes, HEK cells, lipid vesicles) indicating no agonist response to acetylcholine or serotonin (data not shown).

### Conformational changes upon binding of CHAPS

A comparison of the CHAPS-bound structure with the apo state reveals concerted conformational changes. In addition to the local rearrangements of side chains in the ligand-binding site (described above) we observe a clear change in the quaternary conformation of Alpo4 (Figure 3, Supplementary Video 1). Upon binding of CHAPS, the ECD rotates 9-degree clockwise relative to the TMD (when viewed from the extracellular side; Figure 3a, b). The binding of CHAPS is associated with local conformational changes in ECD. First, the tip of the loop C (residues 197-205) shifts by about 3 Å and extends (Figure 3a, d) even though its density in the apo state is somewhat ambiguous (Supplementary Figure 6). The loop movement is accompanied by changes in the orientation of aromatic sidechains (Y195, W197, and Y199) that allow accommodating the zwitterionic moiety of the CHAPS molecule (Figures 2, 3d; Supplementary Video 1). On the complimentary subunit, loop F (residues 171-184) shifts by 3 to 5 Å toward the sterol group of CHAPS. This results in a small (~1 Å) rearrangement of the ECD-TMD linker (Figure 3a, e). The ECD protomers show concerted movements as rigid bodies. They rotate by ~3 degrees around the domain center such that the epical regions of ECD move in the direction of neighboring ECD subunit in a clockwise fashion, whereases TMD-facing ends move counterclockwise (Figure 3a, Supplementary Video 1). The whole ECD assembly rearranges as tightly packed domino tiles.

**Figure 3.**
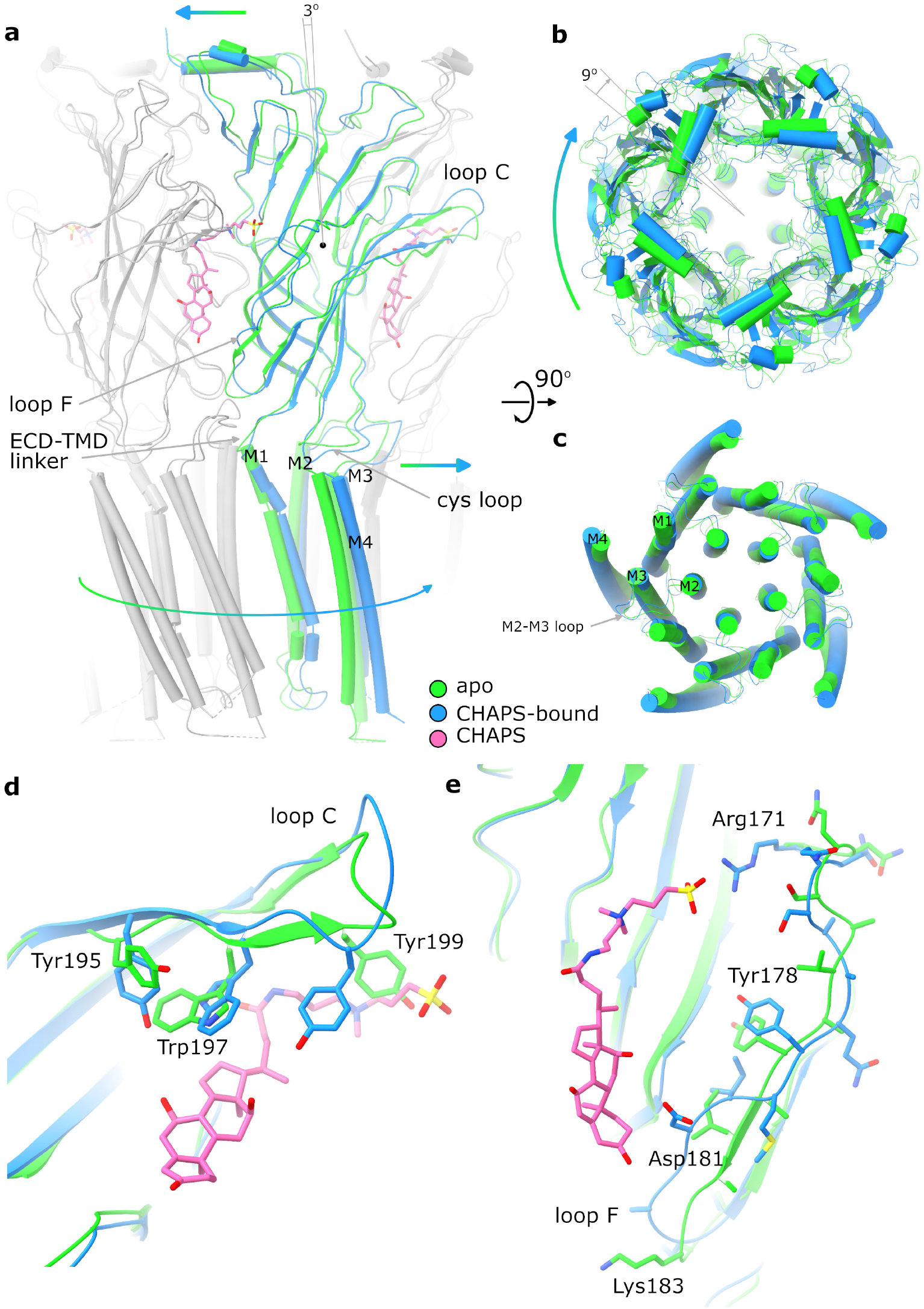
Conformational changes between apo and CHAPS-bound Alpo4. **(a)** Structures of apo (green) and CHAPS-bound (light blue) Alpo4 are overlayed. The ECD was aligned between the structures. Bound CHAPS is shown in pink for reference. Only one subunit from each pentamer is colored, others are shown in grey for clarity. The binding of CHAPS results in an around 3 degrees clockwise rotation. The approximate rotation center is indicated by the black dot. The rotation of the membrane domain is indicated by an arrow. **(b)** Relative rotation of TMD and ECD. The structures are aligned to TMDs. **(c)** Same as panel **b** but only TMDs are shown. Changes in TMD associated with CHAPS binding are minor. **(d, e)** Close-up of conformational changes in loops C and F, respectively.

We can speculate that the combination of CHAPS-induced rigid body tilt of the individual ECDs relative to each other and local rearrangements in the F loop leads to the rotation of TMD relative to ECD. This rotation is accommodated by the bending of the first two helical turns of the M1 helix, between 1 and 3 Å, an extension of the FPF motif on the Cys-loop, and a 3-5 Å shift of M2-M3 loop that follows rigid body movement rotation of the Cys-loop (Figure 3a, c, Supplementary Video 1).

A quaternary twist is known to be associated with gating transitions in characterized pLGICs^10–12,30–33^. The ECD consistently rotates counterclockwise (when viewed from the extracellular side), which contrasts with the clockwise rotation observed in Alpo4 upon binding of CHAPS. The difference in the direction of rotation differentiates general channel gating transitions from the one we observe in Alpo4.

### Structure of the pore domain

In the TMD, all four TM helices are well resolved allowing for unambiguous assignment of the helix register. The densities for the M1-M3 helices are of excellent quality whereas the peripheral M4 displayed higher mobility (Supplementary Figure 6). The ion pore is located along the 5-fold rotational symmetry axis and is formed exclusively by the M2 helices. It has a circa 15 Å long hydrophobic patch in the outer leaflet of the membrane formed by 3 helical turns (residues 9’L, 13’L, and 16’M; Figure 4a). On the extracellular side, the hydrophobic region is flanked by negatively charged aspartate residues, 20’D, whereas on the intracellular side glycines G’6 create a cavity within the pore (Figure 4b) followed by rings of threonines (2’T) and conserved glutamates (−1’E) which usually play a role of the selectivity filter in cation-selective pLGICs^34^. Thus, the charge distribution along the pore is consistent with Alpo4 being selective for cations.

**Figure 4.**
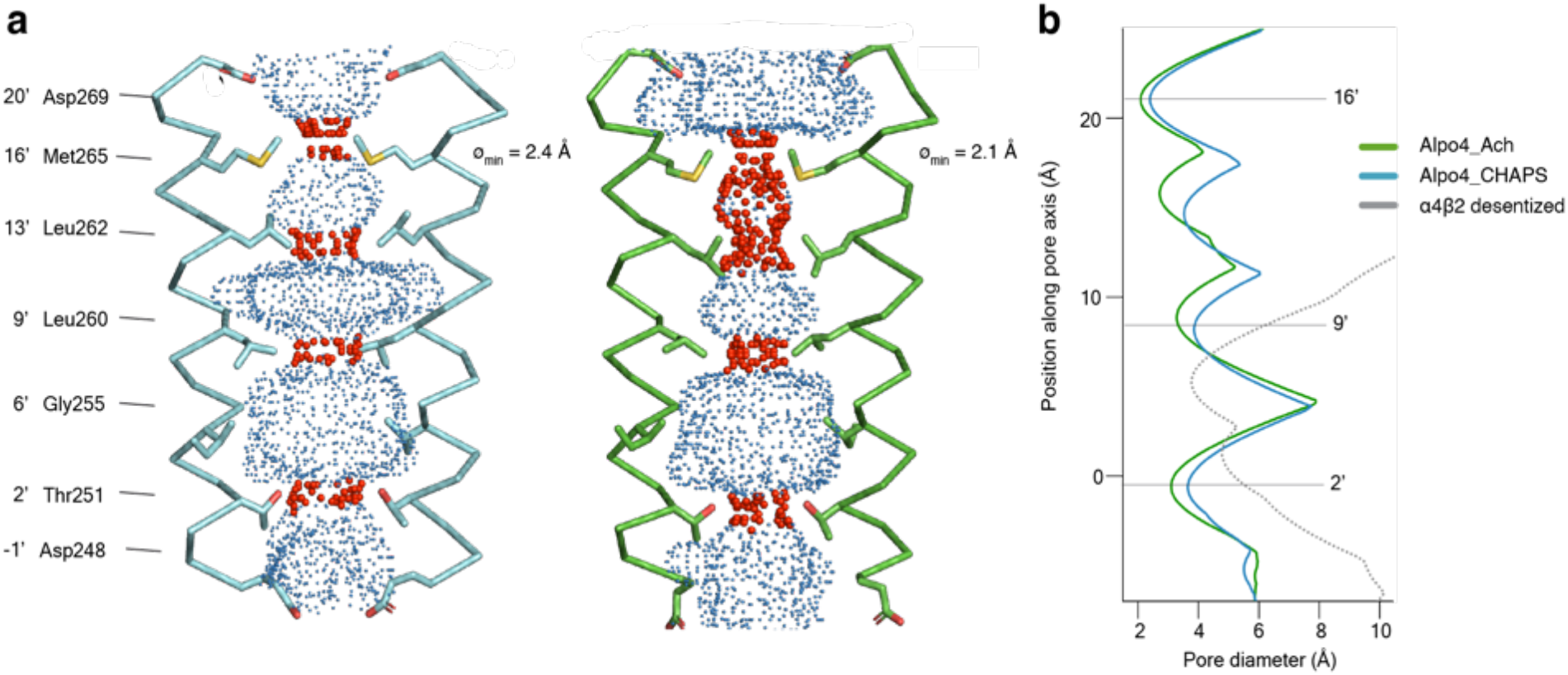
Permeation pathway of Alpo4. **(a)** Pore diameter calculated by HOLE^58^ and represented as dots for Alpo4^CHAPS^ (blue) and Alpo4^APO^ (green). Only M2 is shown in cartoon and the pore-facing residues are shown as sticks. Constrictions are shown in red. **(b)** Pore diameter along the channel axis for Alpo4^CHAPS^ (blue) and Alpo4^APO^ (green). The zero value along the channel axis corresponds to position 2’ (Thr251). The pore profile for the desensitized state of nAChR α4β2 (PDB: 5KXI) (grey) is shown for comparison.

A highly unusual feature in Alpo4 is the presence of bulky M265 residues at the 16’ position. It forms the narrowest and most hydrophobic constriction (diameter of 2.1 Å) that likely functions as a gate. Constriction at this position is absent in other structurally characterized nAChRs but was observed in bacterial Cys-loop receptor homolog ELIC in which the pore is constricted by a Phe residue at the extracellular surface^35^. This observation led us to speculate that the narrow 16’ constriction prevented ion permeation, leading to a lack of agonist responses in our earlier electrophysiological ligand screenings. Therefore, we explored a M16’L mutation, which unfortunately was also unresponsive to acetylcholine or serotonin (data not shown). Next, the M16’L mutation was combined with the well-described L9’T mutation, which slows desensitization, converts certain antagonists into agonists, and increases Ca^2+^ permeability in α7 nAChRs^36^. In our experiments, the double mutant M16’L/L9’T was still unresponsive to acetylcholine or serotonin. Additionally, we constructed chimeras in which the ECD and TMD were swapped with the α7 nAChR, similar to the α7/5-HT_3_ chimera^37^, but these constructs also remained unresponsive. Finally, we also considered that the 16’ methionine residues could confer redox-sensitive channel regulation, similar to the upper gate formed in TRPV2 ion channels^38^. However, we could not detect any channel activity in the presence of oxidizing (H_2_O_2_) or reducing agents (DTT) (data not shown).

In conclusion, Alpo4 has a pore structure consistent with a cation-selective channel, but with an unusually tight constriction at the 16’M position, the role of which remains unclear.

## Discussion

Here we described the first structure of a lophotrochozoan homopentameric Cys-loop receptor, Alpo4. The structure of the nACh-like receptor from extreme thermophile *Alvinella pompejana* was solved in apo form and in complex with CHAPS and provided insight into the architecture of the channel and its unexpected interaction with the sterol derivative.

### A possible function of Alpo4

Recent structural work on channels such as *Torpedo* nAChR, zebrafish GlyRα1, mouse 5-HT_3_, and human α7 nAChR revealed that representative Cys-loop receptors do not share a single conserved gating mechanism, highlighting their functional diversification^10,12,30,33^. In lophotrochozoans, the vast expansion of nAChR genes, their high sequence diversity and spatial expression profiles imply diversified function^39^. Our phylogenetic analysis revealed that Alpo4 doesn’t cluster with homolog lophotrochozoans receptors, except those found in *Capitella teleta*, suggesting a faster or less constrained evolution of the channel. Multiple sequence alignment shows the presence of key aromatic ligand-binding residues and the absence of the characteristic vicinal disulfide in the tip of loop C indicating that Alpo4 lacks signature amino acids of an α-like nAChR subunit^40^. The loss of the CC-tip is also observed in the Alpo3-like polychaetes receptors but not in the other lophotrochozoans organisms (Supplementary figure 1b).

Despite exhaustive screening efforts to identify the ligand, this Cys-loop receptor remains an orphan. Our efforts to identify its ligand included expression in *Xenopus* oocytes followed by two-electrode voltage-clamp electrophysiology with a library of >30 compounds known to act on various Cys-loop receptors^26,36^. As a complementary approach, we expressed Alpo4 in HEK293 cells, followed by patch-clamp electrophysiology with a selected compound library, and employed lipid vesicles to reconstitute detergent-purified Alpo4. The tested compounds, including acetylcholine and serotonin, failed to elicit channel responses (data not shown). Neither could we obtain structural evidence that either acetylcholine or serotonin binds Alpo4 or induces conformational changes in the protein. Alpo4 might be gated by a ligand different from acetylcholine and serotonin consistent with the expansion of nACh genes in lophotrochozoan. Therefore, the agonist may be a chemical that is different from the currently known neurotransmitters in this family of ion channels^26^.

Furthermore, cryo-EM structure-guided mutagenesis of the pore residues and construction of chimeras did not result in an acetylcholine- or serotonin-responsive channel. Additionally, the 16’M pore constriction resembles the formation of an upper gate in TRPV2, which confers redox-sensitive channel regulation^38^. Oxidizing or reducing agents also failed to elicit a response from Alpo4. It is worth noting, that 16’M is not a thermophile-specific residue, as it is also present in the sequences of nAChRs-like proteins from mesophile organisms such as *Pecten maximus* and *Gigantopelta aegis*. It is possible that the channel adapted to extreme environmental conditions and remains closed under laboratory conditions. This however contradicts the electrophysiological experiments that allowed us to identify glycine as the agonist for Alpo5 and Alpo6, which displayed high sequence similarity to glycine receptors (45-50%)^26^, whereas Alpo7 was identified as a pH-gated ion channel^41^. This indicated that thermophilic Alpo Cys-loop receptors can be produced in the functional form in mesophilic expression systems highlighting Alpo4 as an outlier. Among other plausible reasons for the lack of functional activity of Alpo4 in electrophysiological experiments might be a requirement for an accessory β-subunit or a chaperone protein, similar to the recent identification of TMX3 as a co-factor required for the expression of insect nAChRs^42^.

Another unusual feature of the apo structure of Alpo4 is the presence of His131 on loop E, which forms a cation-π interaction with the highly conserved loop B aromatic W159 in the orthosteric binding site. The histidine could potentially coordinate a metal ion or require (de)protonation for the channel to become responsive to an agonist. In the case of CHAPS, this interaction is broken enabling coordination of the quaternary ammonium group by W159. Although CHAPS is not expected to be a native ligand in *Alvinella pompejana*, our structures reveal it interacts with the ECD and induces an atypical quaternary twist movement.

### Bivalent channel modulator

Pharmaceutical α7 nAChR agonists are composed out of three representative groups: a cationic center, a hydrogen bond acceptor, and a hydrophobic element^43^. These pharmacophores are often small molecules that fit in the orthosteric ligandbinding pocket. Here, the example of CHAPS binding creates an unusual precedent for the binding of a channel modulator that interacts with two regions of the ECD, one in the conserved ligand-binding pocket and another with the poorly conserved crevice outside the orthosteric site (Figure 1, Supplementary Figure 4). The sterol group connected by a linker binds in between subunits and induces conformational changes, therefore it likely plays an active role in the observed quaternary twist. This finding hints at a possible strategy for designing specific Cys-loop channel modulators wherein the orthosteric binder is complemented by a chemical group binding at the interface between the subunits. In most channels a crevice is present at the subunit interface (Supplementary Figure 4b), permitting a design of a specific binder. Because the interface is poorly conserved between the nAChRs, the binding of the second group can be designed specifically for a particular channel thereby increasing the specificity of a pharmacologically active molecule.

This study contributes to a better understanding of Cys-loop receptors in lophotrochozoans, highlighting Alpo4 as a member with a yet-to-be-identified neurotransmitter agonist. Our findings that sterol derivatives bind at the orthosteric binding site hint toward new strategies for designing specific channel modulators.

## Material and methods

### Phylogenetic tree construction

A sequence similarity search using BLASTP and the amino acid sequence of Alpo4 as a reference was performed against the genome of *Capitella teleta* (Polychaetes), the closest match to Alpo4, and other annelids with annotated genomes: *Dimorphilus gyrociliatus* (Polychaetes), *Owenia fusiformis* (Paleoannelids), *Hirudo verbana* (Clitellates), *Helobdella robusta* (Clitellates) and proteins of mollusca *Crassostrea virginca* (Bivalvia), *Crassostrea gigas* (Bivalvia), *Mizuhopecten yessoensis* (Bivalvia), *Pecten maximus* (Bivalvia) and *Pomacea canaliculata* (Gastropoda). Sequences with an E-value lower than 1e-5 were used for multiple sequence alignment using CLC v 21.0.5 sequence manager (Qiagen). After multiple rounds of alignments and manual removing non-nAChR sequences a set of 2047 proteins were obtained. To simplify the analysis, only one isoform of each receptor was retained, and a phylogenetic tree was constructed using the CLC sequence manager with the Maximum Likelihood method using 647 sequences. Confidence values were obtained using bootstrapping with 100 sequences. Here, family numbers were generated to identify which nAChRs cluster together as indicated on the phylogenetic tree. For the smaller phylogenetic trees of family 41A (Alpo3-like) and family 43A (Alpo4-like), only 1 isoform for each selected Lophotrochozoan member was used with the addition of *Octopus sensis* (OS), *Gigantopelta aegis* (GA), *Aplysia californica* (AC) and *Pomacea canaliculata* (PC).

### Protein expression and purification

Wild-type Alpo4 was expressed and purified as previously described with some modifications^26^. Briefly, His-tagged Alpo4 was expressed for 72 hours in Sf9 insect cells. Cells were pelleted, flash frozen and stored at −80°C. For protein purification, resuspended cells were lysed through high-pressure homogenization and membranes were isolated after ultracentrifugation. Membranes were solubilized in 10 mM NaPi pH 7.4, 500 mM NaCl, 1% LMNG, and 1% CHAPS for 2 hours at 4°C. The clarified supernatant was incubated with Ni-NTA resin (Roche), extensively washed with 10 mM NaPi pH 7.4, 500 mM NaCl, 0.05% LMNG, and 0.05% CHAPS and Alpo4 was eluted with the wash buffer containing 300 mM imidazole. The purity of the eluted fractions was assessed using SDS-PAGE and the fractions containing Alpo4 were concentrated in 100 kDa molecular weight cutoff concentrators (Invitrogen). Concentrated Alpo4 was injected into a Superose 6 column equilibrated with 25 mM NaPi, 150 mM NaCl, 0.007% CHAPS, 0.003%, LMNG at 4°C. For CHAPS-free Alpo4 preparations (Alpo4^APO_LMNG^, Alpo4^ACH,^ and Alpo4^SER^), CHAPS was omitted from all the buffers used for the solubilization and purification.

### Cryo-EM sample preparation

Graphene oxide coated grids were prepared by incubating 4 μL of 0.9 mg/ml graphene oxide solution for 2 min on R2/1 grids freshly glow discharged for 1 min at 10 mbar pressure and 10 mA current. After blotting with Whatman 1 filter paper, the grids were washed 3 times with H_2_0 and dried for 30 min. Grids with Alpo4^CHAPS^ were prepared by applying 3 μL of Alpo4 (0.04 mg/mL) on the front and 1 μL on the back side of the grid and incubated for 1 min at 100% humidity and 25° C. The sample was plunge-frozen using a CP3 (Gatan) after blotting for 2.7 s from both sides with Whatman 3 filter paper using a blotting force of −1. Grids that produced the Alpo4^APO^ reconstruction were prepared using Alpo4 purified similar to Alpo4^CHAPS^ sample, except that size-exclusion chromatography (SEC) was performed using a buffer without CHAPS. Next, 1 mM acetylcholine was added to the protein solution and incubated for 30 min on ice. A volume of 3 μL protein solution (0.04 mg/mL) was applied to the front of the graphene oxide-coated R2/1 grid and 1 μL of the buffer CHAPS-free SEC buffer was applied on the back of the grid. After 1 min incubation, the grid was blotted and plunge-frozen as described above. The grids with CHAPS-free Alpo4 samples (Alpo^APO_LMNG^, Alpo^ACH^, and Alpo^SER^) were prepared using the graphene oxide-coated grids by applying 2 μL of Alpo4 (0.05 mg/mL) solution on the front and 1 μL of the buffer on the back side of the grid and incubating for 1 min. Acetylcholine (1 mM) or serotonin (1 mM) were added to Alpo4^ACH^, and Alpo4^SER^ samples, respectively, and incubated for 30 min on ice prior to applying the protein solution on the grid. The grids were blotted as described above.

### Data collection

The data were collected on a JEOL CryoARM 300 transmission electron microscope (TEM) equipped with an Omega Filter (20 eV slit) using SerialEM v3.8.0^44^. The Alpo4^CHAPS^ dataset was collected on a Gatan summit K2 direct electron detector operating in counting mode. 5003 movies were collected using a nominal magnification of 60,000 (the calibrated pixel size of 0.782 Å/pixel) using a defocus range of 1.6 to 2.8 μm. Each movie consisted of 50 frames with 0.2 sec/frame exposure and recorded with an electron flux of 3.7 e^-^/sec/Å^2^. The Alpo4^APO^ dataset comprised 7153 movies collected at a nominal magnification of 60,000 (the calibrated pixel size of 0.784 Å/pixel) on K3 direct electron detector using a defocus range of 0.8 to 2.8 μm. Each movie contained 61 frames of 0.038 sec exposure each using a dose of 18.2 e^-^/sec pixels. The Alpo4^APO_LMNG^, Alpo4^ACH^, and Alpo4^SER^ datasets comprised 11,895, 13,850, and 2201 movies, respectively collected at a nominal magnification 60,000 (the calibrated pixel size of 0.760 Å/pixel) using a defocus range of 1.0 to 2.4 μm. Each movie contained 59 frames of 0.05 sec exposure using a dose of 18.2 e-/sec pixels.

### Cryo-EM image processing

For all datasets, movie alignment was done with MotionCorr2^45^ and contrast transfer function (CTF) was estimated with CtfFind4^46^. For the Alpo4^CHAPS^ dataset, particles were picked with crYOLO v1.8^47^, extracted in a box size of 368 pixels, and decimated 4 times (3.1 Å/pixel) followed by two rounds of 2D classification in Relion 3.0^48^ with between 20 and 50 classes. An *ab initio* model was generated without imposing symmetry and used as a starting model after low pass filtering to a resolution of 15 Å. Multiple rounds of 3D classification were performed with 3 to 6 classes without applying symmetry in Relion 3.0. The best-resolved classes were auto-refined in Relion imposing C5 symmetry. The selected particles were re-extracted without binning and subjected to Bayesian polishing and per-particle defocus refinement followed by a final 3D auto-refinement step^49^. During the postprocessing step, a soft mask was applied. Local resolution was estimated using Relion 3.0.

In the case of the Alpo4^APO^ dataset, images were denoised using the JANNI software^50^ prior to particle picking using crYOLO. Particles were extracted in a box size of 358 pixels and imported into CryoSPARC v3.3.1^51^ and 5 rounds of reference-free 2D classification were performed. Five *ab initio* models were generated with C1 symmetry and a maximum resolution of 12 Å. Subsequent hetero- and non-uniform refinement with C5 symmetry and a dynamic mask resulted in a model with a resolution of 3.4 Å. Both global and local CTF refinement was performed in CryoSPARC. The particles were used for a final *ab initio* model calculation followed by non-uniform refinement with C5 symmetry. This resulted in a 3D reconstruction at a resolution of 3.4 Å. Similar processing strategies were applied to the Alpo^APO_LMNG^, Alpo^ACH^, and Alpo^SER^ datasets and resulted in reconstructions at a resolution between 4.1 and 6.1 Å (Supplementary Table 1, Supplementary Figs 3, 5, 7, 9-11).

### Cryo-EM model building and structure analysis

The model of mouse serotonin 5-HT_3_ receptor (PDB 6HIQ^29^) was used as an initial model for building the structure of Alpo4^CHAPS^. A monomer was fitted into the cryo-EM map (JiggleFit) and manually rebuild using Coot 0.9.5^52^. The model was expanded to a pentamer by applying C5 symmetry operator in Phenix 1.19^53^. The resulting model was refined in Phenix using real_space_refinement routine^54^. Here, global minimization, rigid body fit, and local rotamer fitting were performed with C5 symmetry imposed. After each refinement cycle, the model was manually adjusted in Coot. Due to the poor density of helix M4, it was initially built as a polyalanine chain. Later, the register of the M4 helix was determined from the 3.4 Å Alpo4^APO^ reconstruction and fitted into Alpo4^CHAPS^ map. The refined Alpo4^APO^ model was placed in Alpo^APO_LMNG^, Alpo^ACH^, and Alpo^SER^ reconstructions as a rigid body and refined using the real_space_refinement routine. The models were validated using MolProbity^55^, Phenix^53^ and Coot. Figures were generated using UCSF Chimera and UCSF ChimeraX v1.3^56^, and PyMol v2.4.0. Structure-based sequence alignment was performed in PROMALS3D^57^.

## Data Availability

The cryo-EM density maps and atomic models generated in this study have been deposited in the PDB and EMDB database under accession codes: 8BYI / EMDB-16326 (Alpo4^CHAPS^), 8BXF / EMDB-16317 (Alpo4^APO^), 8BX5 / EMDB-16308 (Alpo4^LMNG_APO^), 8BXB / EMDB-16314 (Alpo4^ACH^), 8BKE / EMDB-16316 (Alpo4^COMB^), 8BKD / EMDB-16315 (Alpo4^SER^).

The atomic models used in this study are available in the PDB database under accession code 6NP0 (5-HT3), 5KXI (nAChα4), 5KXI (nACh β2), 4HFI (GLIC), and 6HJX (ELIC).

## Acknowledgment

We are indebted to Dr. Adam Schröfel and Dr. Marcus Fislage for their assistance with cryo-EM data collection at BECM. We thank Marijke Brams and Mieke Nys for assistance with baculovirus production and protein expression. We would like to acknowledge the funding provided by Vlaams Instituut voor Biotechnologie, Fonds Wetenschappelijk Onderzoek (Grant Nos. G0H5916N, G054617N to R.G.E., and G0H5916N to S.D.G). C. U. was supported by grants from FWO-Vlaanderen (G0C1319N, G087921N) and KU Leuven (C3/19/023, C14/17/093).

