## Supplementary Information for "Sterol derivative binding to the orthosteric site causes conformational changes in an invertebrate Cys-loop receptor"

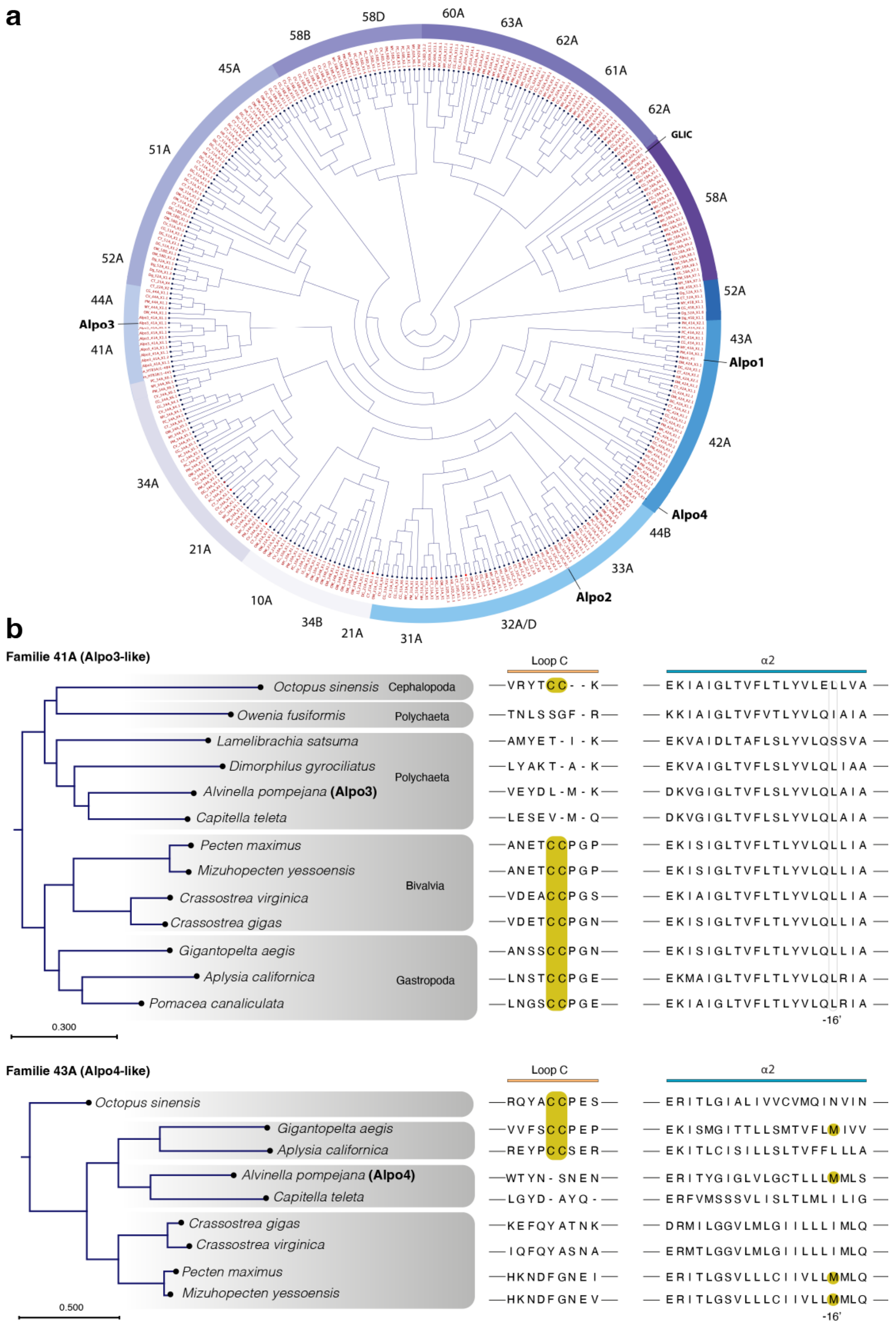


**Supplementary Figure 1 Phylogenetic analysis of nAChR in lophotrochozoans. (a)** Maximum likelihood phylogeny of lophotrochozoans nAChR receptors using one isoform of each protein sequence from genomes of the following annelids: *Capitella teleta (*CT)*, Dimorphilus gyrociliatus* (DM), *Owenia fusiformis* (OW), *Hirudo verbana* (HV), *Helobdella robusta* (HR) and proteins of Mollusca *Crassostrea virginca* (MV), *Crassostrea gigas* (CG), *Mizuhopecten yessoensis* (MY), *Pecten maximus* (PM) and *Pomacea canaliculata* (PC). Based on this classification, the nAChR were grouped in families as indicated in the tree. Positions of Alpo1-4 are indicated. (b) Phylogenetic tree and multiple sequence alignment of proteins sequences corresponding to family 41A (Alpo3-like) and 43A (Alpo4-like). Only one isoform for each selected Lophotrochozoan member was used with the addition of *Octopus sensis* (OS), *Gigantopelta aegis* (GA), *Aplysia californica* (AC), and *Pomacea canaliculata* (PC). The alignments show the conservation of the characteristic CC tip (yellow) in the ligand-binding pocket. Alignment of the M2 sequences reveals proteins with methionine on position 16'.

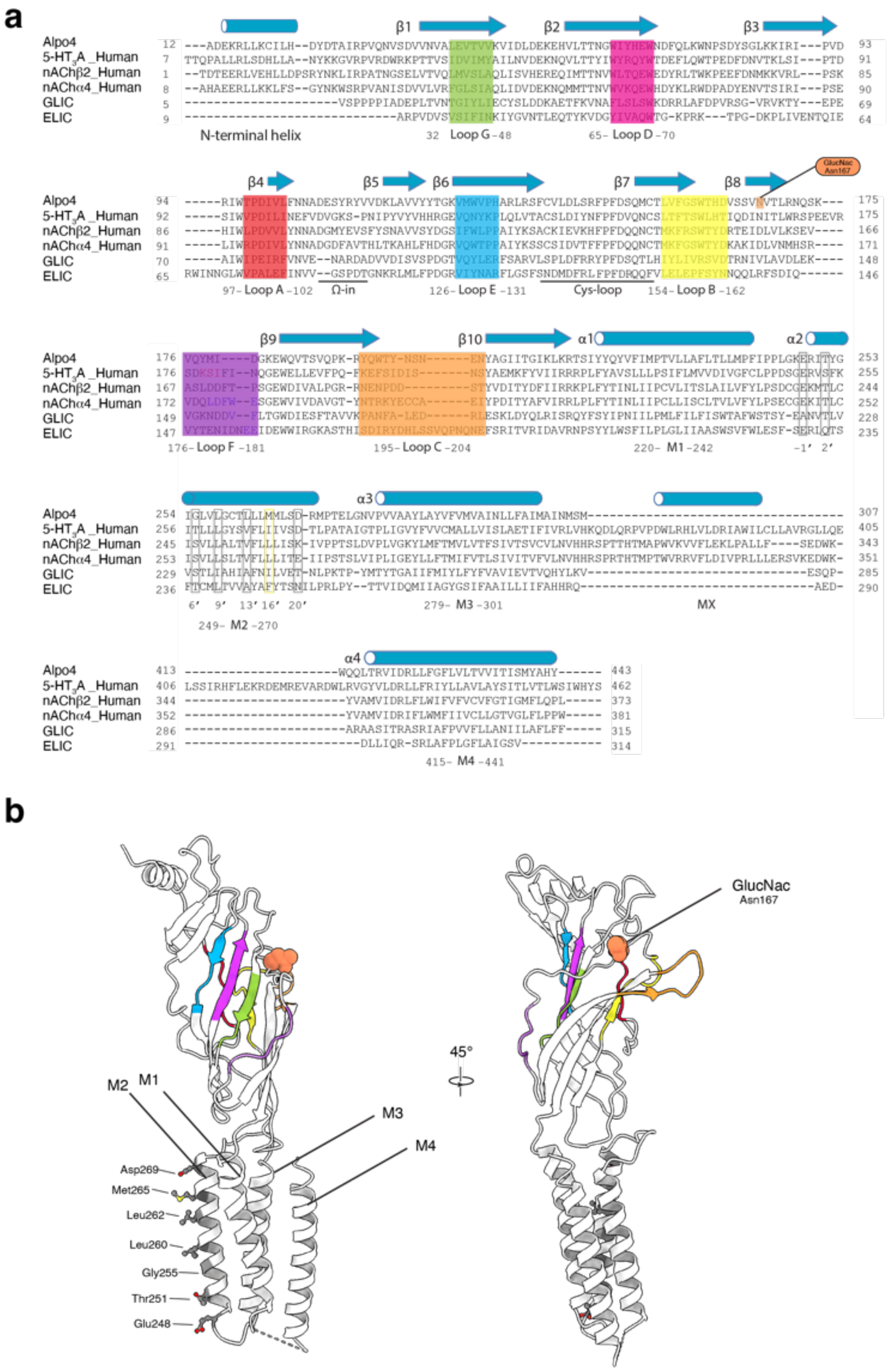


**Supplementary Figure 2 Structure-based sequence alignment of cation-selective pLGICs. (a)** Sequence alignment of selected pLGIC by PROMALS3D. The following structural models were used for the alignment: Alpo4$^{ACH}$ (chain A), 5-HT$_3$A (PDB: 6NPO, chain A), nACh α4 (PDB: 5KXI, chain A), nACh β2 (PDB: 5KXI, chain B), GLIC (PDB: 4HFI, chain A), and ELIC (PDB: 6HJX, chain A). Secondary structure elements are indicated. Loops A-F are colored and pore-facing residues are outlined by grey rectangles. One glycosylation site at Asn167 was identified in the structure of Alpo4. ISD, residues 308-412, are not resolved in Alpo4 maps. The sequence numbering of structural elements corresponds to Alpo4. **(b)** The structure of the Alpo4 monomer is shown in cartoon representation. Loops A-F are color-coded as in panel a, GlucNac on Asn167 is shown in coral, and the pore-facing residues are shown in stick and ball representation.

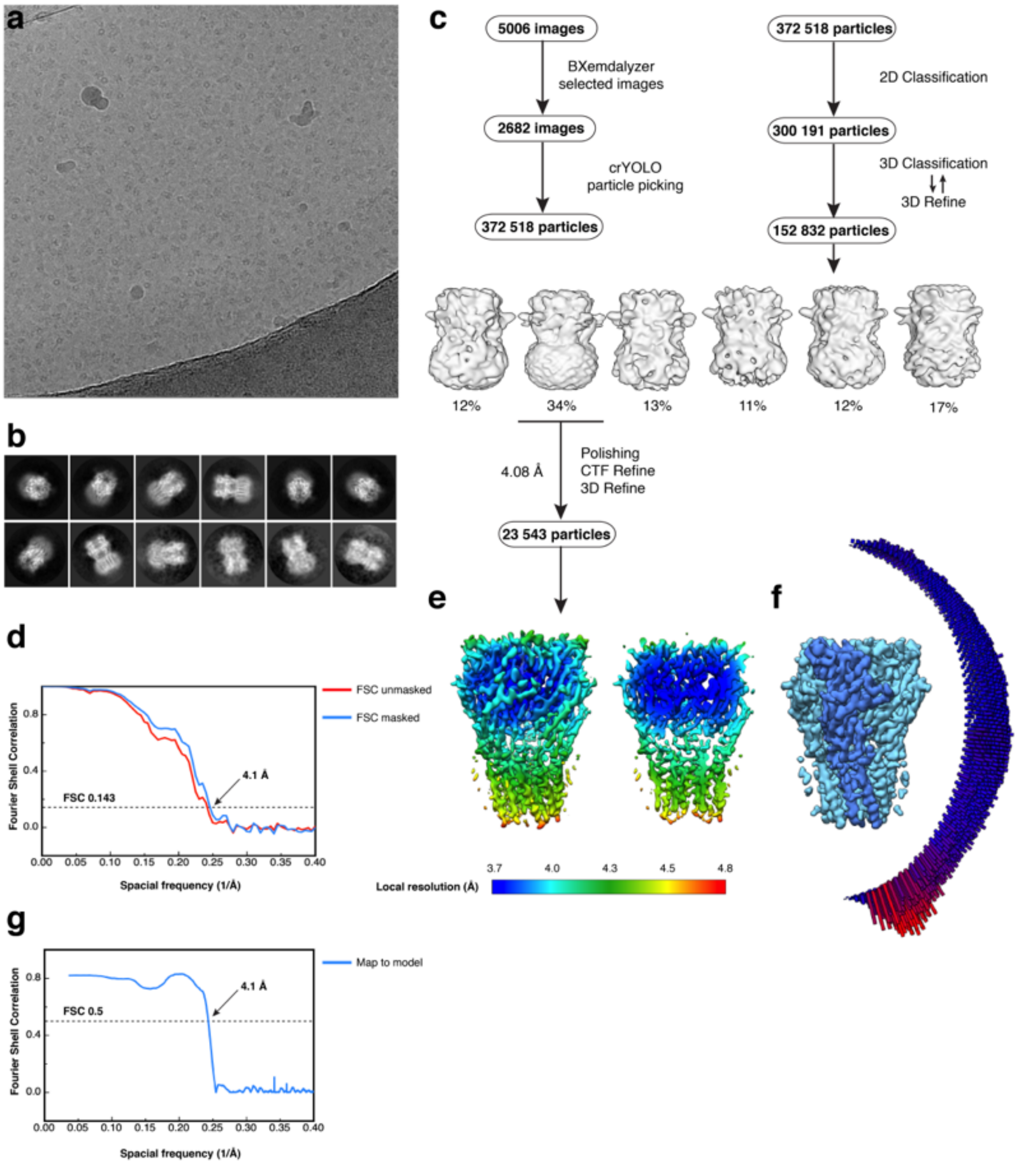


**Supplementary Figure 3 Electron microscopy, classification, and 3D reconstruction for the Alpo4$^{CHAPS}$ dataset**. **(a)** Representative micrograph of Alpo4$^{CHAPS}$ embedded in vitreous ice on graphene oxide coated R2/1 grids. **(b)** The 2D class averages. **(c)** Image selection, particle picking, classification, and refinement workflow. **(d)** Gold-standard Fourier Shell Correlation (FSC) curves are shown for unmasked (red) and masked (blue) reconstructions. The horizontal dashed line indicates the 0.143 cutoff threshold. **(e)** Side-view and 'clipped' view of the final masked reconstruction colored by local resolution calculated using Relion 3.0. **(f)** Angular distribution of particle orientations for the final reconstruction. **(g)** Model-to-map FSC indicates a nominal resolution of 4.1 Å using the 0.5 FSC threshold.

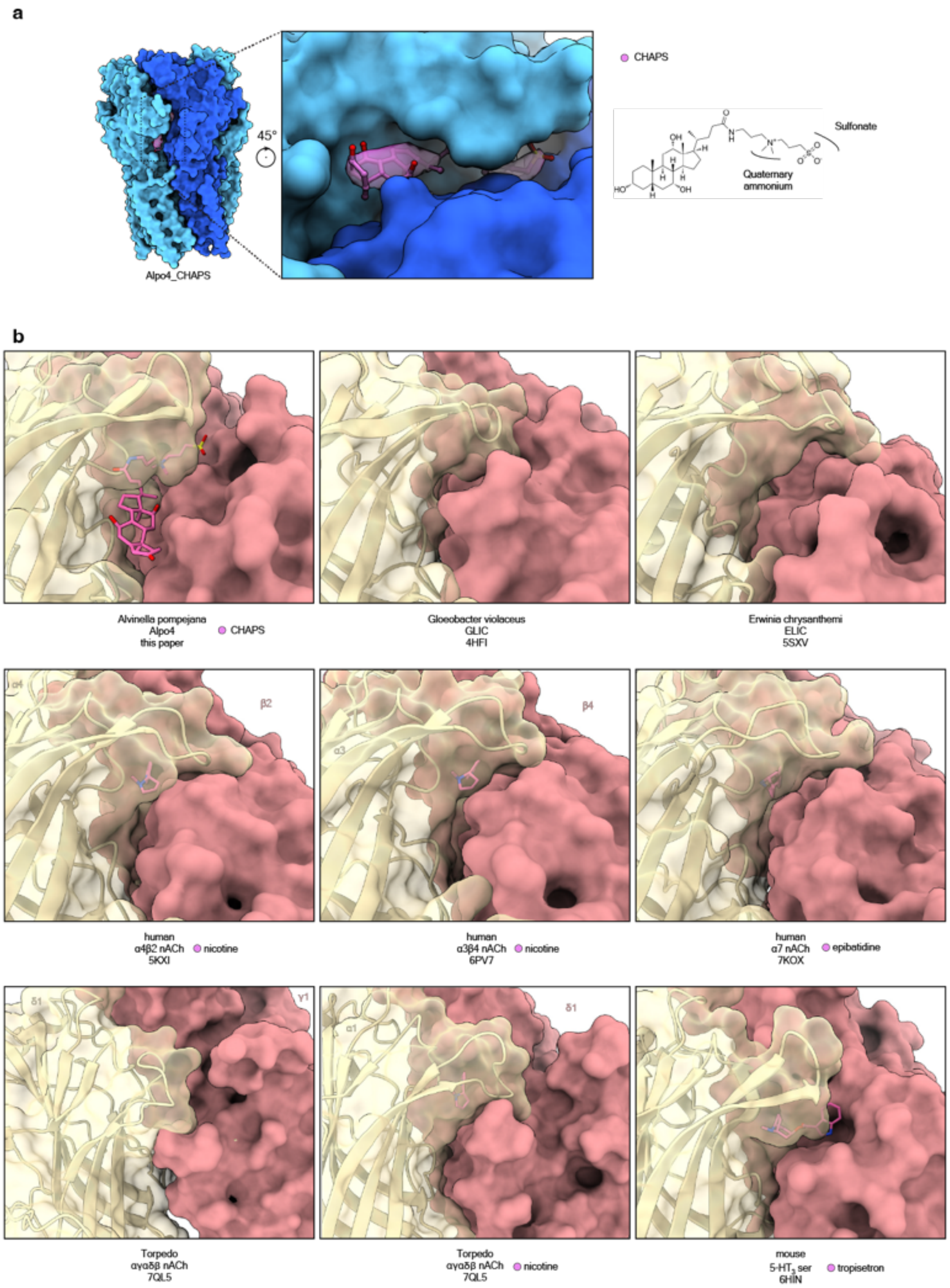


**Supplementary Figure 4 Sterol-binding pocket is specific for Alpo4. (a)** Surface representation of Alpo4$^{CHAPS}$ in blue highlights the position of additional density found in the cryo-EM map (violet). The modeled CHAPS molecule is shown in ball and sticks representation. **(b)** Ligand binding pockets of different pGLIC reveals accessible area. Ligands are shown in pink.

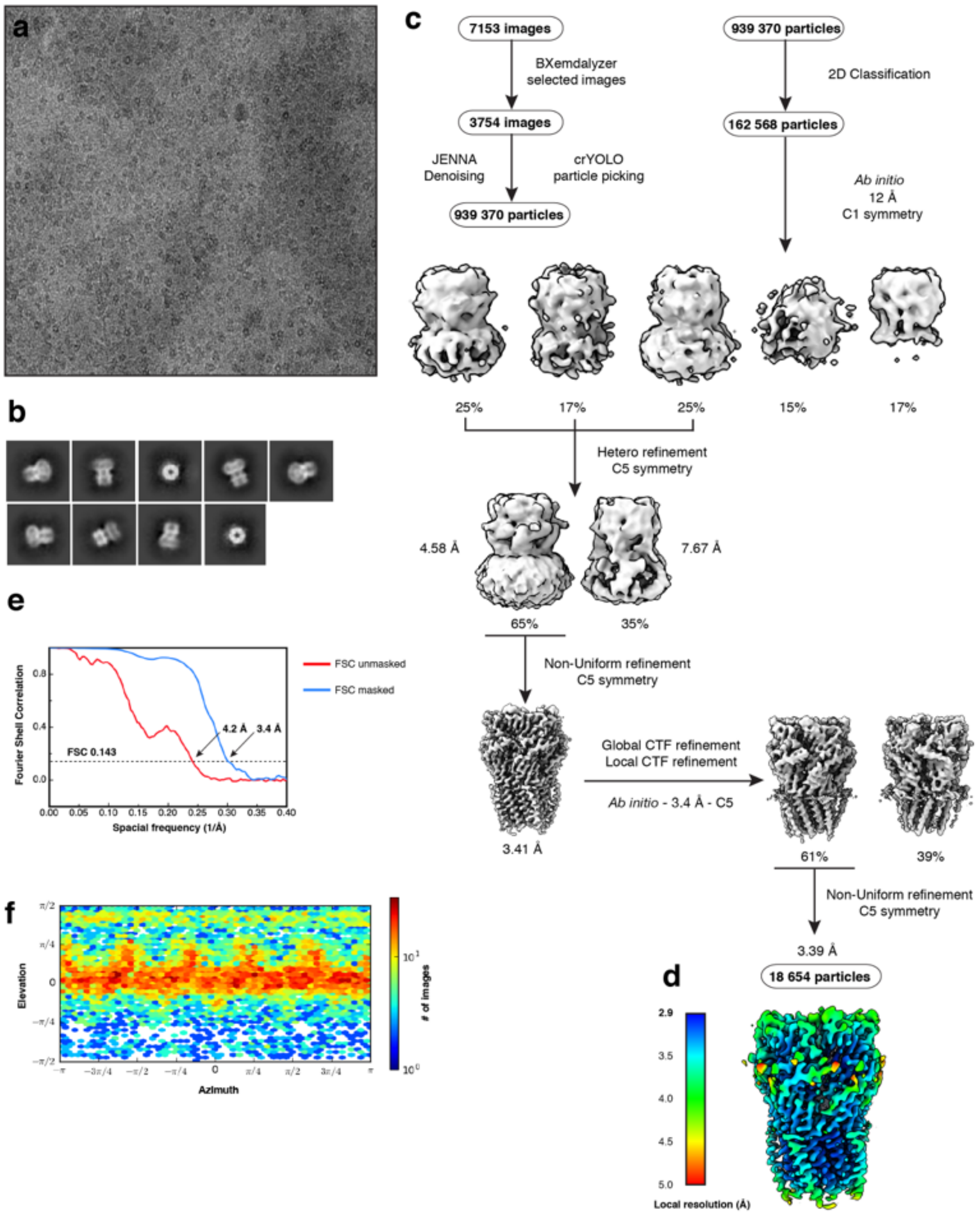


**Supplementary Figure 5 Electron microscopy, classification, and 3D reconstruction for the Alpo4$^{APO}$ dataset. (a)** Representative micrograph of Alpo4$^{APO}$ embedded in vitreous ice on graphene oxide coated R2/1 grids. **(b)** The 2D class averages. **(c)** Image selection, particle picking, classification, and refinement workflow in cryoSPARC. **(d)** The local resolution of the final reconstruction is calculated using cryoSPARC. **(e)** Gold-standard Fourier Shell Correlation (FSC) curves are shown for unmasked (red) and masked (blue) reconstructions. The horizontal dashed line indicates the 0.143 cutoff threshold. **(f)** Heat map of the angular distribution of the final reconstruction as calculated in cryoSPARC.

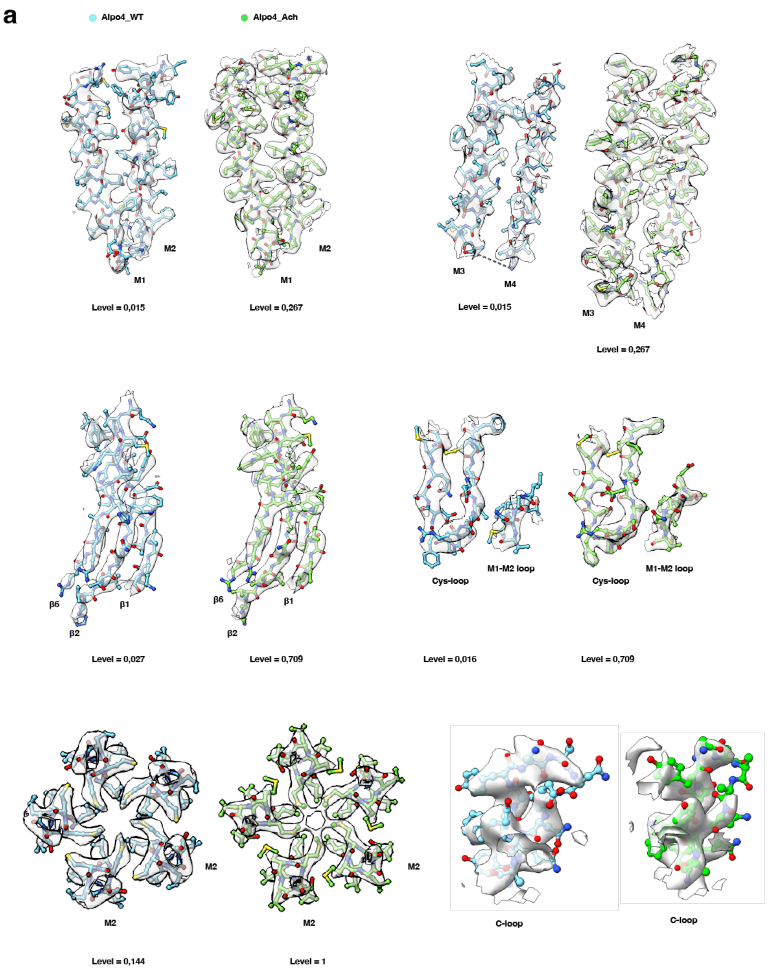


**Supplementary Figure 6 Quality of the cryo-EM maps.** The cryo-EM maps and models are shown for representative regions of Alpo4$^{CHAPS}$ (blue) and Alpo4$^{APO}$ (green).

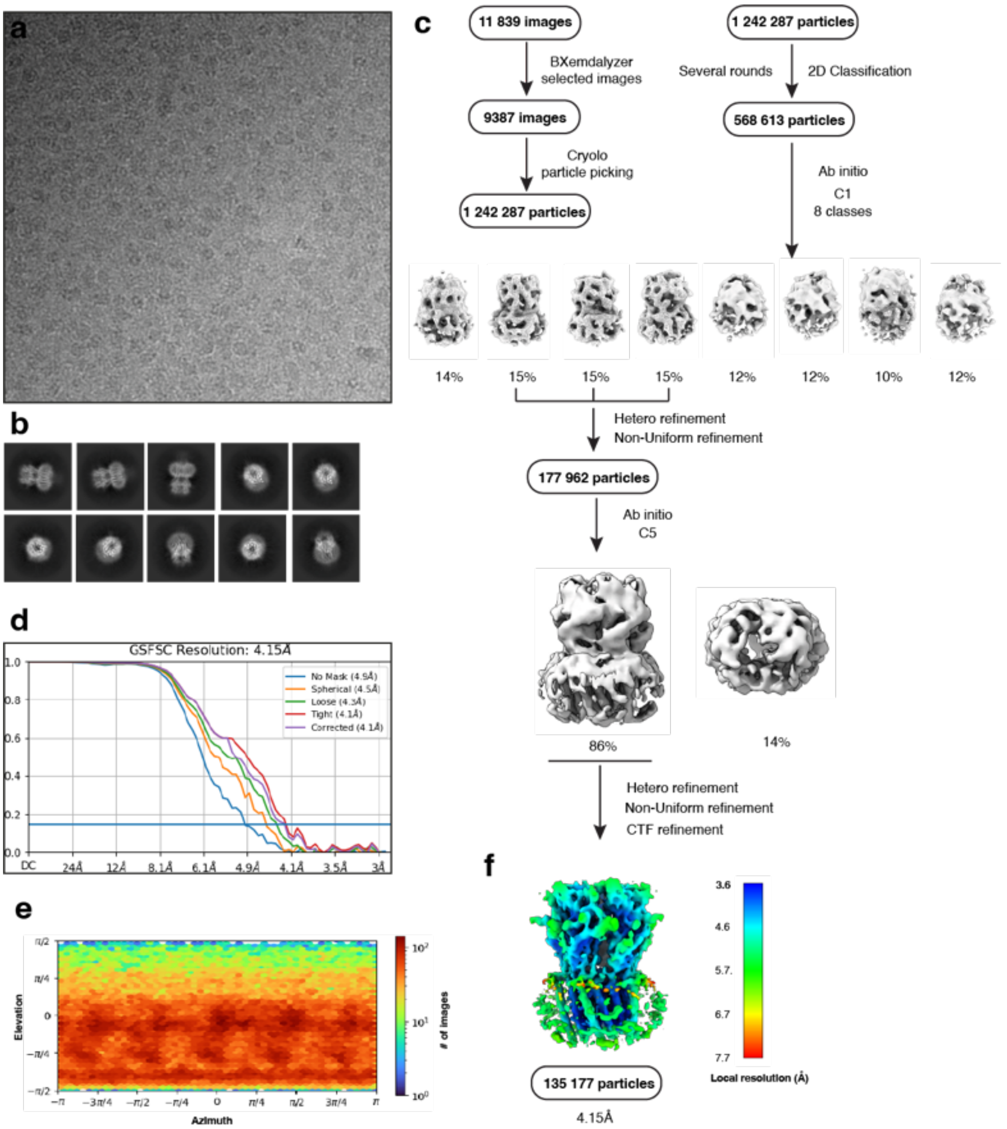


**Supplementary figure 7 Electron microscopy, classification, and 3D reconstruction for the Alpo4$^{APO_LMNG}$ (LMNG only) dataset**. **(a)** Representative micrograph of Alpo4$^{APO_LMNG}$ embedded in vitreous ice on graphene oxide coated R2/1 grids and **(b)** the resulting 2D class averages of the final model. **(c)** Image selection, particle picking, classification, and refinement workflow in cryoSPARC. **(d)** Gold-standard Fourier shell correlation (FSC) curves with the horizontal dashed lines highlighting the 0.143 threshold. The unmasked reconstruction (red) reaches 4.9 Å while the tight dynamic masked reconstruction (blue) reaches 4.1 Å. **(e)** Heat map of the angular distribution of the final reconstruction as calculated in cryoSPARC. **(f)** The local resolution of the final reconstruction is calculated using cryoSPARC.

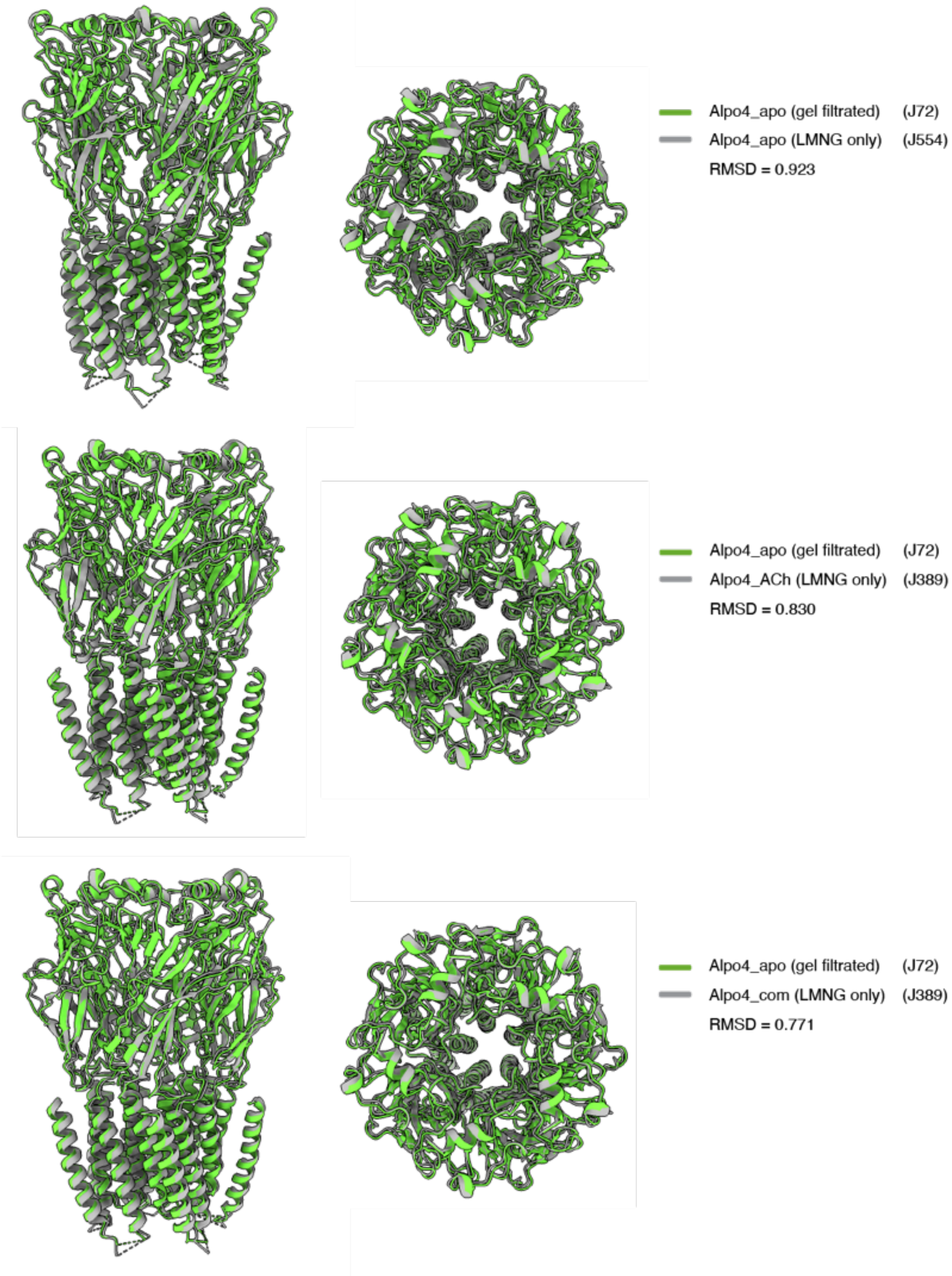


**Supplementary Figure 8 Superposition of Alpo4 structures.** Alpo4$^{APO}$ (gel filtrated) is shown in green and superimposed on Alpo4$^{APO_LMNG}$, Alpo4$^{ACH,}$ and Alpo4$^{COMBINED}$ (LMNG only).

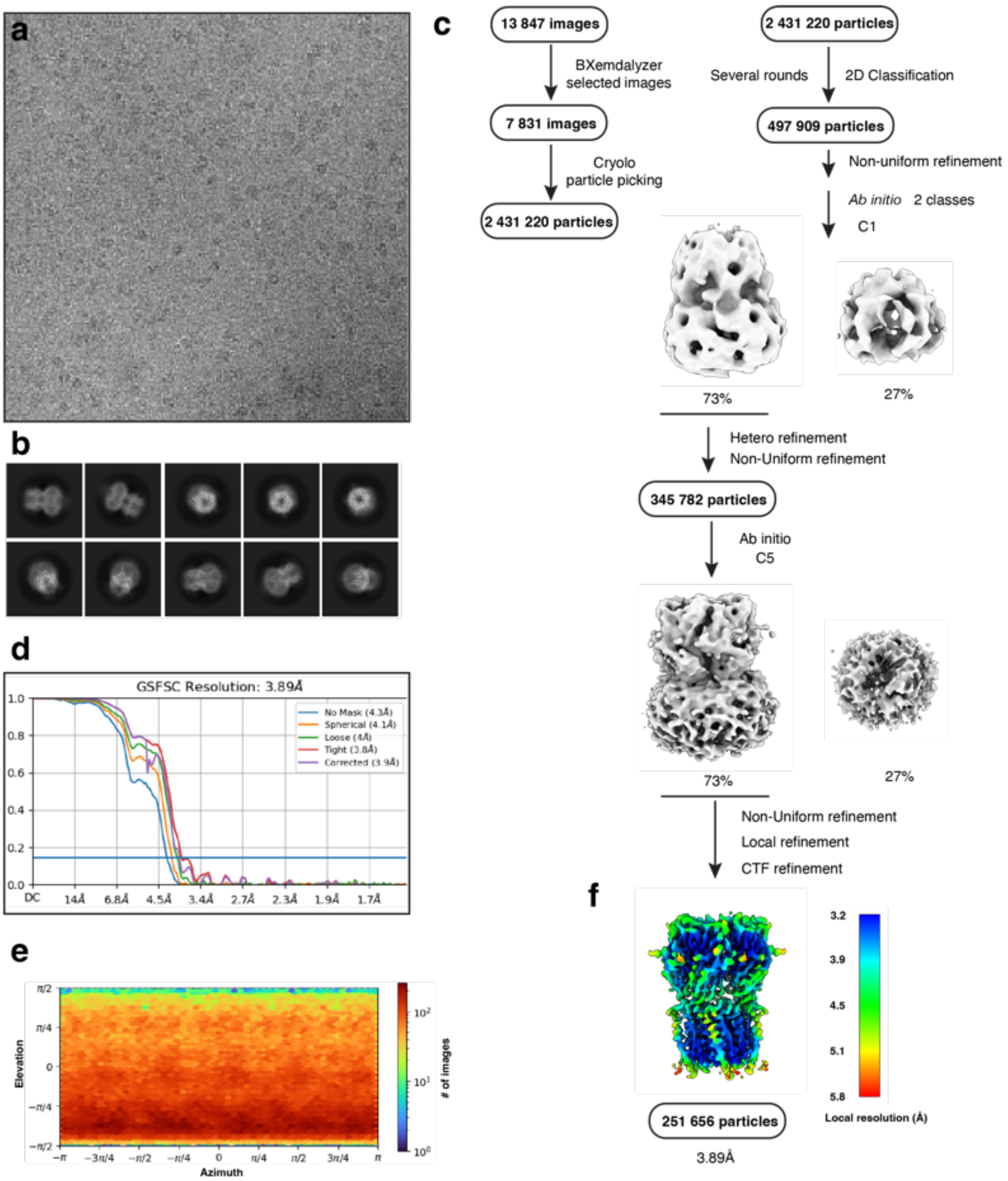


**Supplementary figure 9 Electron microscopy, classification, and 3D reconstruction for the Alpo4$^{ACH}$ (LMNG only) dataset**. **(a)** Representative micrograph of Alpo4$^{ACH}$ embedded in vitreous ice on graphene oxide coated R2/1 grids and **(b)** the resulting 2D class averages of the final model. **(c)** Image selection, particle picking, classification, and refinement workflow in cryoSPARC. **(d)** Gold-standard Fourier shell correlation (FSC) curves with the horizontal dashed lines highlighting the 0.143 threshold. The unmasked reconstruction (red) reaches 4.3 Å while the tight dynamic masked reconstruction (blue) reaches 3.9 Å. **(e)** Heat map of the angular distribution of the final reconstruction as calculated in cryoSPARC. **(f)** The local resolution of the final reconstruction is calculated using cryoSPARC.

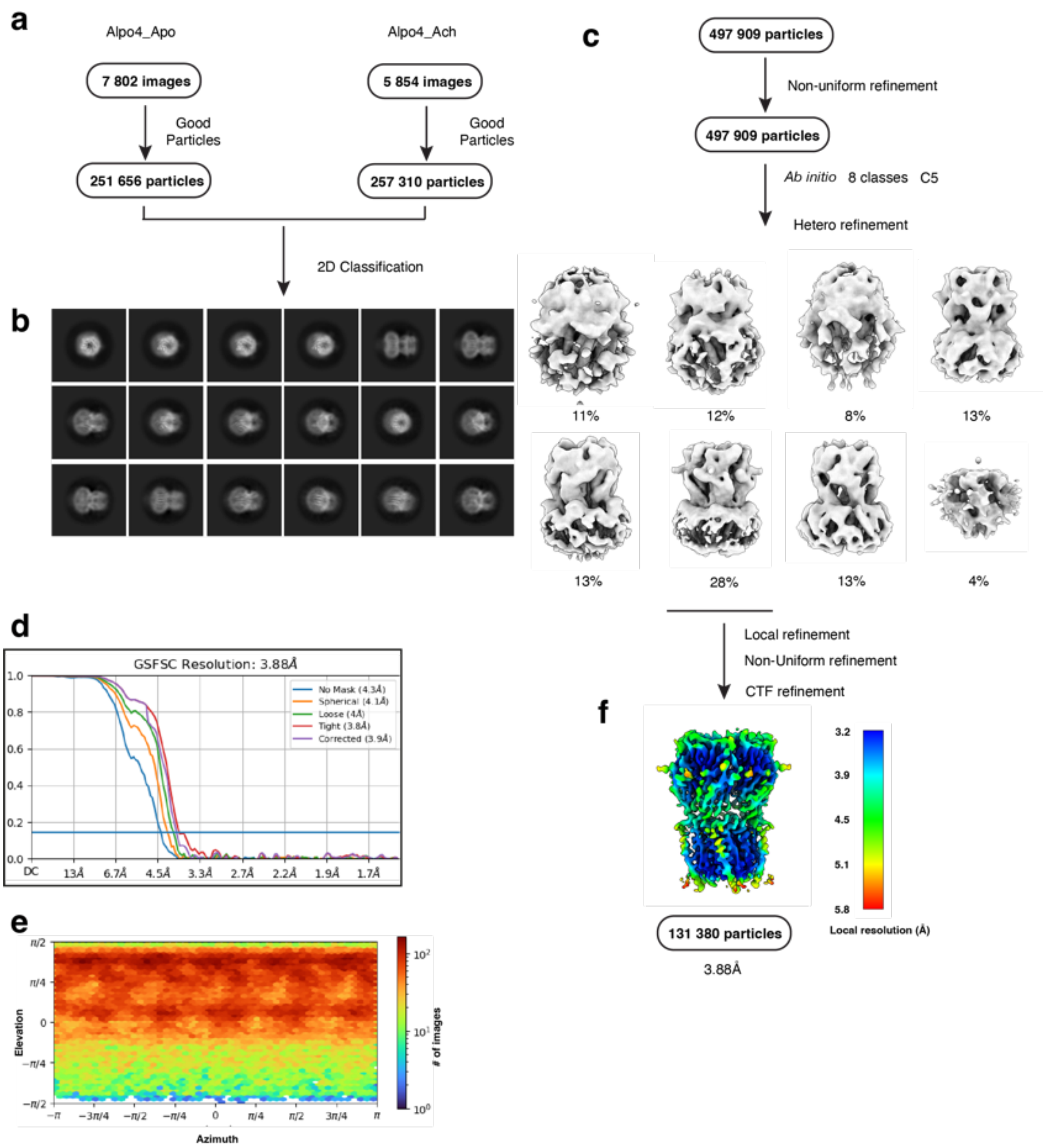


**Supplementary figure 10 Electron microscopy, classification, and 3D reconstruction for the Alpo4$^{combined}$ (LMNG only) dataset. (a)** Cryo-EM workflow of combining Alpo4$^{APO_LMNG}$ and Alpo4$^{ACh}$ particles in cryoSPARC. **(b)** representative 2D classifications. **(c)** refinement workflow. **(d)** Gold-standard Fourier shell correlation (FSC) curves with the horizontal dashed lines highlighting the 0.143 threshold. The unmasked reconstruction (red) reaches 4.3 Å while the tight dynamic masked reconstruction (blue) reaches 3.9 Å. **(e)** Heat map of the angular distribution of the final reconstruction as calculated in cryoSPARC. **(f)** The local resolution of the final reconstruction is calculated using cryoSPARC.

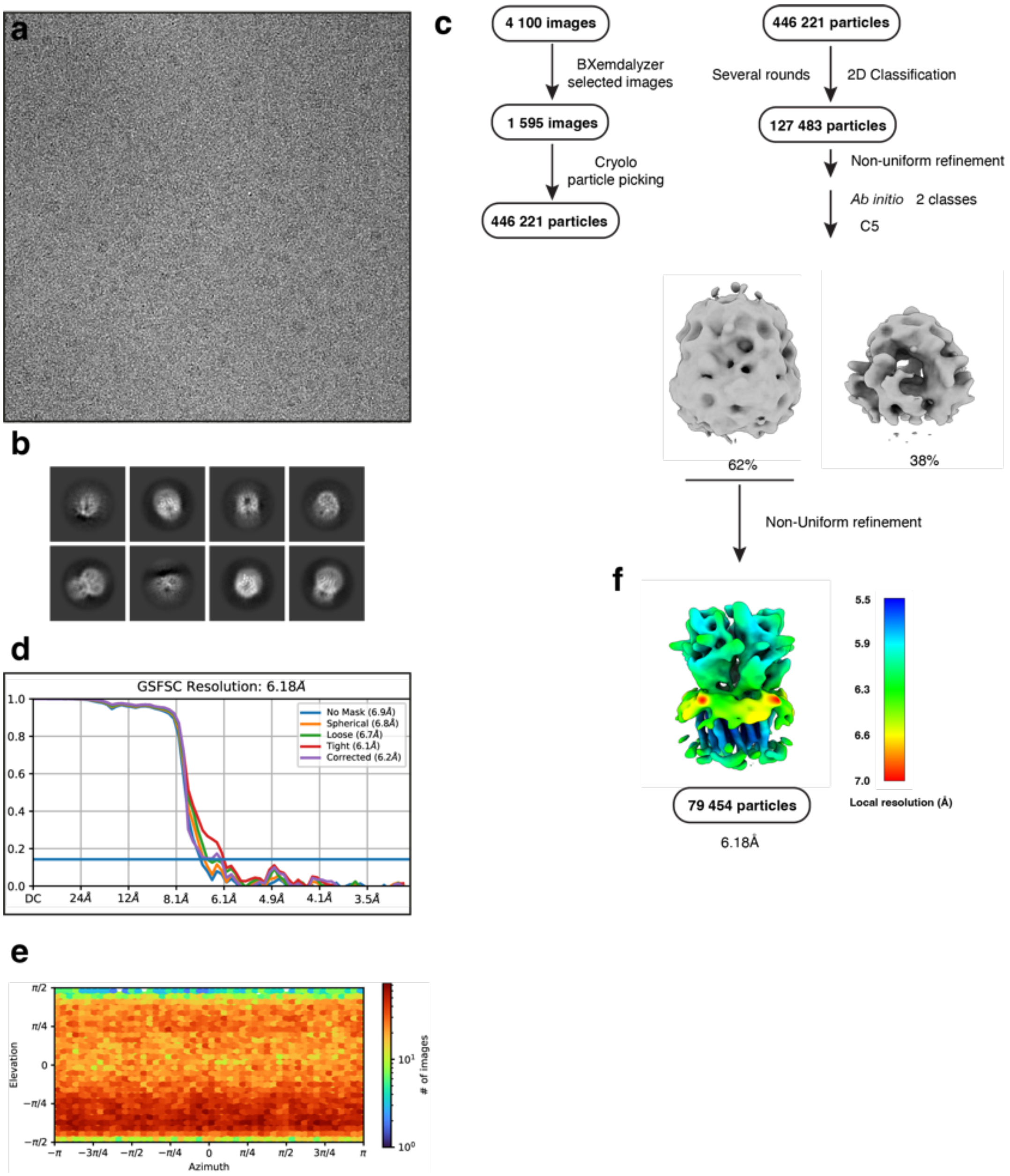


**Supplementary figure 11 Electron microscopy, classification, and 3D reconstruction for the Alpo4$^{SER}$ (LMNG only) dataset. (a)** Representative micrograph of Alpo4$^{SER}$ embedded in vitreous ice on graphene oxide coated R2/1 grids and **(b)** the resulting 2D class averages of the final model. **(c)** Image selection, particle picking, classification, and refinement workflow in cryoSPARC. **(d)** Gold-standard Fourier shell correlation (FSC) curves with the horizontal dashed lines highlighting the 0.143 threshold. The unmasked reconstruction (red) reaches 6.9 Å while the tight dynamic masked reconstruction (blue) reaches 6.2 Å. **(e)** Heat map of the angular distribution of the final reconstruction as calculated in cryoSPARC. **(f)** The local resolution of the final reconstruction is calculated using cryoSPARC.

### Tables

#### Table 1 Statistics of cryo-EM data collection, data processing, and model refinement

| **Data deposition** | | | | | | |
|---|---|---|---|---|---|---|
| Alpo4 ID: | Alpo4$^{CHAPS}$ | Alpo4$^{APO}$ | Alpo4$^{APO_LMNG}$ | Alpo4$^{ACH}$ | Alpo4$^{COMB}$ | Alpo4$^{SER}$ |
| PDB ID: | 8BYI | 8BXF | 8BX5 | 8BXB | 8BXE | 8BXD |
| EMDB ID: | 16326 | 16317 | 16308 | 16314 | 16316 | 16315 |
| **Data collection** | | | | | | |
| Microscope | JOEL CRYOARM300 | | | | | |
| Acceleration voltage [kV] | 300 | | | | | |
| Energy filter | In-column Omega energy filter | | | | | |
| Energy filter slit width [eV] | 20 | | | | | |
| Spherical aberration [mm] | 2.55 | | | | | |
| Magnification | 60 000 | | | | | |
| Detector | Gatan K2 | Gatan K3 | Gatan K3 | Gatan K3 | Gatan K3 | Gatan K3 |
| Refined pixel size [Å] | 0.782 | 0.784 | 0.7596 | 0.7596 | 0.7596 | 0.7596 |
| Exposure time [s] | | | 2.985 | 3.955 | 2.985/3.955 | 3.955 |
| Number of frames | 61 | 61 | 59 | 59 | 59 | 59 |
| Electron exposure [e-/Å$^2$] | 37 | 30 | 45 | 59 | 45/59 | 59 |
| Defocus range [µm] | 1.6 - 2.8 | 0.8 - 2.8 | 1.0 - 2.4 | 1.0 - 2.4 | 1.0 - 2.4 | 1.0 - 2.4 |
| Collected images | 5003 | 7153 | 11 839 | 13 830 | 25 669 | 2200 |
| Used images | 2682 | 3754 | 9387 | 7816 | 13 647 | 1595 |
| Particles picked | 372 518 | 939 370 | 1 242 287 | 2 597 373 | 508 966 | 446 221 |
| **Data processing** | | | | | | |
| Symmetry | C5 | C5 | C5 | C5 | C5 | C5 |
| Particles refined | 23 543 | 18 654 | 135 177 | 251 656 | 131 380 | 79 454 |
| Final resolution [Å], FSC=0.143 | 4.1 | 3.4 | 4.2 | 3.9 | 3.9 | 6.2 |
| Sharpening B-factor [Å$^2$] | -176 | -139 | -297 | -170 | -235 | -1062 |
| Local resolution range [Å] | | 2.9 - 5.0 | | | | |
| **Model refinement** | | | | | | |
| Refinement package | PHENIX 1.19 | | | | | |
| Initial model used | 6HIQ | Alpo4$^{CHAPS}$ | Alpo4$^{ACH}$ | Alpo4$^{APO}$ | Alpo4$^{ACH}$ | Alpo4$^{ACH}$ |
| Model resolution [Å$^2$], FSC=0.5 | 4.2 | 3.9 | 4.5 | 4.2 | 4.1 | |
| Model composition | | | | | | |
| Non-hydrogen protein atoms | 13 321 | 12 692 | 12 550 | 12 547 | 12 553 | |
| Protein residues | 1600 | 1630 | 1630 | 1630 | 1600 | |
| Ligands | CPS/NAG: 5 | NAG: 5 | NAG: 5 | NAG: 5 | NAG: 5 | |
| B-factors mean [Å$^2$] | | | | | | |
| Protein | 63 | 93 | 203 | 54 | 93 | |
| Ligand | 40 | 108 | 199 | 54 | 111 | |
| R.M.S deviations | | | | | | |
| Bond lengths (Å) | 0.002 | 0.005 | 0.003 | 0.004 | 0.004 | |
| Bond angles (°) | 0.688 | 1.240 | 0.736 | 0.779 | 1.055 | |
| Validation | | | | | | |
| Molprobity score | 1.9 | 1.6 | 1.8 | 2.3 | 1.9 | |
| Clashscore | 24.1 | 4.8 | 14.1 | 7.2 | 12.5 | |
| Poor rotamers (%) | 0 | 0.4 | 0 | 5.5 | 0 | |
| Ramachandran plot | | | | | | |
| Favored (%) | 98 | 96 | 97 | 96 | 95 | |
| Allowed (%) | 2 | 4 | 3 | 4 | 5 | |
| Disallowed (%) | 0 | 0 | 0 | 0 | 0 | |

*Combined J554 and J329 dataset

**Supplementary Table 1 Channel parameter values for anionic pentameric ligand-gated ion channels.** Ion channel PDB files, channel conformation, ligands, and narrowest constriction residues are shown.

| Cationic | PDB | State | Minimal diameter (Å) | Constriction Residue | Position | Ligand | Detergent/ND | Lipids |
|---|---|---|---|---|---|---|---|---|
| Alpo4 | | Closed | ~2.4 | Met265 | 16' | / | LMNG | |
| | | Desensitized | ~2.1 | Met265 | 16' | CHAPS | LMNG/CHAPS | |
| | 6HIN | Open | ~6.0 | Ser253 | 2' | Serotonin | Detergent | |
| mouse 5-HT3A | 6HIO | Pre-wet | ~4.4 | Leu260 | 9' | Serotonin | Detergent | |
| | 6HIQ | Desensitized | ~5.0 | Ser253 | 2' | TMPPAA | Detergent | |
| | 6HIS | Closed | ~4.4 | Glu250 | -1' | Tropisetron | Detergent | |
| human α4β2 | 5KXI_A | Desensitized | ~3.8 | Glu | -1' | Nicotine | Detergent | |
| | 5KXI_B | | | | | | | |
| | 6PV7_A | Desensitized | ~1.7 | Glu | -1' | Nicotine | Saposin A | |
| human α3β4 | 6PV7_B | | | | | | | |
| | 6PV8_A | Desensitized | ~1.7 | Glu | -1' | AT-1001 | DDM | |
| | 6PV8_B | | | | | | | |
| human α7 | 7KOO | Resting | ~2.4 | Leu247 | 9' | α-bungarotoxin | | 10 uM SBL / 0.25% Cholesterol |
| | 7KOX | Activated | ~7.2 | | | Epi + PNU | 1 mM DDM | |
| | 7KOQ | Desensitized | ~4.3 | Glu237 | -1' | Epibatidine | | |
| | 7QKO | Resting | ~2.0 | Leu258 | 16' | Apo | | |
| | 7QL6 | Desensitized | ~6.0 | Thr/Ser244 | 2' | Carbachol | MSP2N2 | NaCholate / SBL |
| | 7QL5 | | | | | Nicotine | | |
| torpedo αγαδβ | 7SMM | Resting | ~2.5 | Leu258 | 16' | Apo | | |
| | 7SMQ | | | | | Apo / chol | | |
| | 7SMR | | | | | Carbachol | Saposin A | Cholesterol / SBL (1:4) |
| | 7SMS | Desensitized | ~4.5 | Thr/Ser244 | 2' | d-Turbo | | |
| | 7SMT | | | | | d-Turbo / carb | | |
| | 6HJX | Closed | ~1.3 | Phe247 | 16' | / | UDM | |
| | 8D64 | Unassigned | <2.0 | | | Cysteamine | | POPC |
| ELIC | 8D63 | Resting | <1.0 | Phe247 | 16' | Apo | | |
| | 8D66 | Unassigned | <2.0 | | | Cysteamine | MSP1E3D1 | 2POPC |
| | 8D65 | Resting | <1.0 | | | Apo | | 1POPE |
| ELIC3 | 8D67 | Unassigned | <2.0 | Leu240 | 9' | Cysteamine | | 1POPG |
| ELIC5 | 8D68 | Open | ~12.6 | / | / | Cysteamine | | |
| | 8F32 | Pre-active | | Phe247 | 16' | | SMA | |
| ELIC | 8733 | Pre-active | | | | Cysteamine | Saposin A | 2POPC |
| | 8F34 | Desensitized | ~3.0 | | 2' | | spMSP1D1 | 1POPE |
| | 8F35 | Closed | | | 16' | Apo | spMSP1D1 | 1POPG |
| GLIC | 4HFI | | | Thr226 | 2' | DDM | | |

**Supplementary Movie 1** Transition of Alpo4$^{Apo}$ to Alpo4$^{CHAPS}$ viewed from the extracellular side, side view, the C-loop, and the F-loop.
